# Accessibility of the Shine–Dalgarno sequence dictates N-terminal codon bias in *E. coli*

**DOI:** 10.1101/195727

**Authors:** Sanchari Bhattacharyya, William M. Jacobs, Bharat V. Adkar, Jin Yan, Wenli Zhang, Eugene I. Shakhnovich

## Abstract

Despite considerable efforts, no physical mechanism has been shown to explain N-terminal codon bias in prokaryotic genomes. Using a systematic study of synonymous substitutions in two endogenous *E. coli* genes, we show that interactions between the coding region and the upstream Shine–Dalgarno (SD) sequence modulate the efficiency of translation initiation, affecting both intracellular mRNA and protein levels due to the inherent coupling of transcription and translation in *E. coli*. We further demonstrate that far-downstream mutations can also modulate mRNA levels by occluding the SD sequence through the formation of non-equilibrium secondary structures. By contrast, a non-endogenous RNA polymerase that decouples transcription and translation largely alleviates the effects of synonymous substitutions on mRNA levels. Finally, a complementary statistical analysis of the *E. coli* genome specifically implicates avoidance of intra-molecular base-pairing with the SD sequence. Our results provide general physical insights into the coding-level features that optimize protein expression in prokaryotes.

## Introduction

The genetic code is degenerate, such that almost all amino acids are coded by more than one codon. However, on a genome-wide scale, not all synonymous codons are used with the same frequency. This codon-usage bias varies across different organisms (Grantham et al., 1980a; Grantham et al., 1980b) and is reflected in the overall GC content of a genome (Lynn et al., 2002; Muto and Osawa, 1987). Yet within a particular genome, synonymous codons are not distributed randomly. For example, codons that are under-represented in highly expressed genes tend to be translated more slowly and with slightly less accuracy (Bulmer, 1991; Eyre-Walker and Bulmer, 1993; Sharp and Li, 1987), suggesting that essential genes have been optimized for translational efficiency. Even within a gene, rare codons often appear in conserved clusters (Chaney and Clark, 2015; Chaney et al., 2017; Clarke and Clark, 2010; Jacobs and Shakhnovich, 2017), indicating that, in some cases, rare codons might also be under positive selection.

In most prokaryotic and eukaryotic genes, the first ~30 amino acids are more likely than the rest of the protein sequence to be encoded by rare codons (Eyre-Walker and Bulmer, 1993; Tuller et al., 2010). The origin of this ‘N-terminal codon bias’ is debatable, with various mechanisms put forward in different studies. One hypothesis is that rare codons have been selected to reduce the speed of translation at the beginning of genes (Tuller et al., 2010). Such a “ramp” in elongation might reduce the likelihood of ribosomal traffic jams along a transcript, preventing premature termination. However, other studies have implicated RNA secondary structure near the 5’-end of mRNA transcripts. Based on synthetic libraries of 154 synonymous variants of GFP (Kudla et al., 2009) and more than 14,000 gene fragments fused to GFP (Goodman et al., 2013), the folding free energy of truncated mRNA transcripts was found to be the best predictor of the rate of protein production. In these studies, codon rarity did not appear to have a measurable effect on protein expression. Although this proposal has also been supported by a statistical analysis of several bacterial genomes (Bentele et al., 2013), which showed that the presence of rare codons correlates with the formation of weaker mRNA secondary structures in GC-rich genomes, a contradictory experimental study of synonymous variants of two human genes expressed in *E. coli* (Welch et al., 2009) found no statistically significant correlation between protein expression and mRNA folding stability.

A key limitation of these previous experimental studies is their reliance on non-endogenous genes. For example, biases in the selection of promoters and N-terminal regions in the synthetic libraries might give rise to spurious correlations while concealing more fundamental physical reasons for the variability of expression among synonymous variants. The use of libraries with substitutions confined to the 5’-end also hinders our ability to distinguish between competing mechanistic proposals. For these reasons, an experimental study investigating one mutation at a time is needed to understand the physical mechanisms responsible for N-terminal codon bias.

In this paper, we report a systematic study of synonymous substitutions in two endogenous *E. coli* genes: *folA*, which codes for the low-copy essential protein Dihydrofolate Reductase (DHFR; ~40 copies/cell), and *adk*, which codes for the more abundant essential protein Adenylate Kinase (ADK; ~700 copies/cell) (Taniguchi et al., 2010). We find that codon selection near the N-terminus is the primary determinant of both protein expression and the steady-state mRNA levels in *E. coli*, with the first rare codon appearing to play a crucial role in both *folA* and *adk*. Our analysis shows that the influence of synonymous substitutions near the N-terminus is best explained by the folding stability of secondary structures near the 5’-end of the mRNA transcript, as opposed to either codon rarity or GC content. In particular, we demonstrate that mutations that lower the intracellular mRNA levels promote the formation of mRNA secondary structures that occlude the upstream Shine–Dalgarno (SD) sequence, thereby affecting the efficiency of translation initiation. Because *E. coli* RNA polymerase tends to carry out transcription cotranslationally (Byrne et al., 1964; Kohler et al., 2017; Miller et al., 1970; Proshkin et al., 2010), the absence of ribosomes on a nascent mRNA chain reduces both the intracellular mRNA and protein levels, presumably due to premature abortion of transcription by Rho (Adhya and Gottesman, 1978; Richardson, 1991). We therefore find that the use of T7 RNA polymerase, which decouples transcription and translation, removes most of the effects of codon optimization on mRNA levels while retaining the effects on protein levels. To highlight the crucial role of the SD sequence in translation initiation, we also show that far-downstream mutations that result in complementary “anti-SD” sequences can modulate mRNA levels as well, although to a lesser extent than substitutions near the N-terminus. Such downstream effects cannot be predicted using equilibrium mRNA folding calculations, but rather require attention to non-equilibrium secondary structures that may form during co-transcriptional mRNA folding. Finally, a complementary statistical analysis of the *E. coli* genome reveals that genes have evolved to minimize binding between coding regions and their respective upstream SD sequences. The SD-occlusion mechanism that we describe therefore appears to be a universal and essential determinant of synonymous codon bias in *E. coli*.

## Results

### Synonymous mutations affect both mRNA and protein levels in *E. coli*

We initially introduced synonymous substitutions directly on the chromosomal copy of *folA* in *E. coli* in order to observe any fitness and proteomic effects in their endogenous context. First, we designed a fully optimized synonymous variant (‘MutAll’) of the *folA* gene, in which all codons were replaced by the most common synonymous codon in the wild-type (WT) sequence (Figure 1A). Contrary to our expectations, we observed a substantial decrease in the exponential-phase growth rate of the *E. coli* population and an increase in the lag time, defined as the time required to achieve the maximum growth rate (Adkar et al., 2017) (Figure 1B). We subsequently determined that these mutations cause a 20-fold decrease in the amount of intra-cellular soluble protein, measured by quantitative Western blot, and a 4-fold drop in intra-cellular mRNA levels, measured via quantitative PCR (qPCR) (Figure 1C). These measurements indicate that, for this gene, codon “optimization” in fact decreases cellular fitness by reducing DHFR expression.

**Figure 1:**
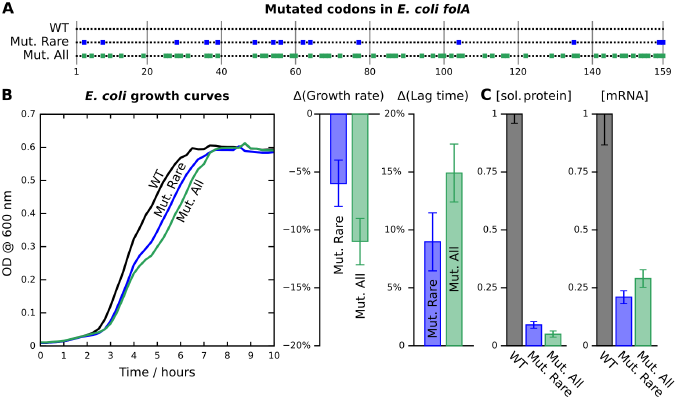
Synonymous substitutions in the chromosomal copy of *E. coli folA* affect cellular fitness, soluble protein abundance, and mRNA levels. (A) The locations of optimized codons within each *folA* construct are indicated by colored squares. (B) Replacing WT codons by their most frequently used synonymous variants on the chromosomal copy of *folA* gene has a deleterious effect on *E. coli* growth, reducing the exponential growth rate and increasing the lag time. (C) Synonymous substitutions affect both the soluble DHFR abundance (measured by quantitative Western blot) and steady-state mRNA transcript levels (measured by qPCR). All values are normalized to chromosomal WT levels.

We further determined that the decreases in soluble-protein and mRNA levels can be attributed almost entirely to the optimization of the 15 “rare” codons, i.e., those codons that occur substantially less frequently than their synonymous alternatives in the most abundant *E. coli* genes (‘MutRare’; Figure 1A,C). This observation is surprising in light of several recent studies, which found that the effects on mRNA abundances due to the introduction of rare codons in *E. coli* (Boel et al., 2016), yeast (Presnyak et al., 2015) and a filamentous fungus (Tokuoka et al., 2008) are typically either deleterious or neutral. Nevertheless, the greatly reduced fitness of the MutRare construct in terms of both growth rate and lag times (Figure 1B) strongly suggests that the presence of rare codons in the WT *folA* gene is not simply a result of neutral drift.

### Systematic codon replacements point to a crucial role of the first rare codon

We next set out to dissect the effects of individual rare codons on the mRNA levels of *folA* and *adk*. In order to perform a systematic investigation, each gene was cloned under an arabinose-inducible pBAD promoter, which allows modulation of expression from 10 to 1000-fold over endogenous levels (Bhattacharyya et al., 2016). Using a fixed arabinose concentration of 0. 05%, we observed 13-fold and 9-fold reductions, respectively, in the mRNA abundances of the MutAll and MutRare *folA* constructs on the plasmid (Figure 2A). In an attempt to further narrow down the search, we tried optimizing only the eight rare codons that are also present in the *folA* gene of the bacterium *Citrobacter koseri*, which shares 96% sequence similarity with the *E. coli folA* gene (‘MutSelect’). MutSelect also showed an 8-fold drop in mRNA levels, suggesting that this subset of rare codons is most critical. The majority of this effect could, in turn, be attributed to substitutions of the first four rare codons (‘MutN-term’) at positions 3, 8, 28, and 36. Finally, we determined that optimization of the first rare codon at position 3 (‘3AGC’) has the single greatest contribution to the decrease in mRNA levels. Maintaining only the first four rare codons while optimizing the remaining rare positions (‘MutC-term’) kept the mRNA abundance close to the WT level. At the same time re-introduction of the WT rare codon at position 3 on the MutRare background (‘MutRare+3AGT’ in *folA*) resulted in mRNA levels that are close to WT.

**Figure 2:**
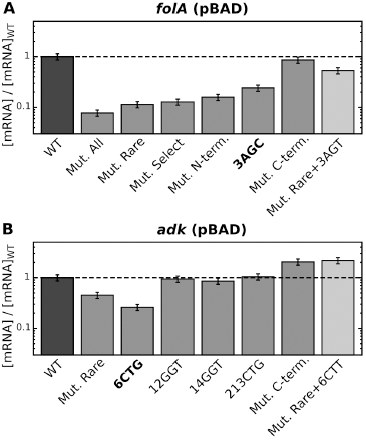
The first rare codon has the largest effect on intracellular mRNA levels. Using an arabinose-inducible pBAD promoter, intra-cellular mRNA levels at an inducer concentration of 0. 05% were measured using quantitative PCR and compared to their WT level (see Table S1). A systematic optimization of rare codons in both (A) *folA* and (B) *adk* revealed that optimizing the first rare codon in each gene has a dominant effect. At the same time, re-introduction of the first rare codon (3AGT for *folA* and 6CTT for *adk*) on the background of the MutRare construct largely rescued the mRNA levels.

To determine if these trends are generalizable beyond *folA*, we carried out a parallel investigation of the *E. coli* gene *adk*. Again, optimization of all six WT rare codons (‘MutRare’) profoundly lowered the mRNA levels, with the dominant contribution coming from the first rare codon at position 6 of *adk* (‘6CTG’ in Figure 2B). Consistent with our findings in *folA*, mutations in the other conserved rare codons of *adk*, at positions 12, 14 and 213, did not contribute to the decrease in mRNA levels. As in the case of *folA*, maintaining the first few rare codons at the N-terminus (Mut C-term) and re-introduction of the WT rare codon at position 6 on the background of ‘MutRare’ (‘MutRare+6CTT’) resulted in mRNA levels close to the WT level. Nevertheless, while the rare codons nearest to the N-terminus were found to modulate the mRNA abundances most dramatically in both genes, some downstream mutations did result in substantial changes (Table S1). These surprising effects will be discussed in detail below.

### The effect of the first rare codon is not due to rarity

Having observed that the greatest mRNA-level effects arise from the first rare codon, we investigated whether the rarity of this codon plays any role. To this end, we mutated position 3 of *folA* to all six Serine codons and position 6 of *adk* to all six Leucine codons on the pBAD plasmid (Figure 3A,B). We found that, in the case of *folA*, only one codon (TCA) maintained the WT mRNA levels, while in the case of *adk*, two codons (CTA and CTC) actually increased the mRNA abundance above the WT level. All other choices resulted in substantial reductions in the mRNA levels. Notably, these effects do not correlate with codon rarity. Meanwhile, the effects of optimizing these key codons were measurably altered by synonymous substitutions at adjacent positions (the second and fourth codons of *folA* and the fifth codon of *adk*), emphasizing that the effects of synonymous substitutions are dependent on their local sequence contexts.

**Figure 3:**
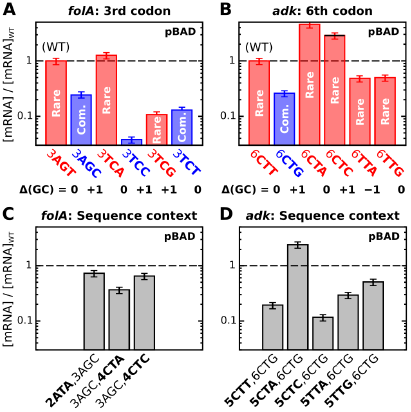
The pivotal role of the first rare codon is not due to rarity. Using an arabinose-inducible pBAD promoter, intra-cellular mRNA levels at an inducer concentration of 0. 05% were measured using quantitative PCR and compared to their WT level (see Table S1). (A) All Serine codons were incorporated at position 3 of *folA*. Other than the WT codon (AGT), only TCA was tolerated. (B) Similarly, all Leucine codons were incorporated at position 6 of *adk*. CTA and CTC were found to increase the mRNA levels relative to the WT. In both panels (A) and (B), blue bars indicate common (optimized) codons while red indicates rare codons. Changes in the GC content, Δ(GC), due to these replacements at position 3 of *folA* and position 6 of *adk* are shown below the construct labels in panels (A) and (B). These constructs demonstrate that neither codon rarity nor GC-content is responsible for the observed effects. (C,D) Substitutions at neighboring positions may rescue the effects of optimizing position 3 of *folA* or position 6 of *adk*, indicating that the role of the codons at these positions is context-dependent.

### Variations in mRNA abundances correlate with the mRNA folding stability

We next sought to determine whether changes in the measured mRNA levels correlate with the predicted conformational stabilities of the mRNA transcripts. After ruling out variations in the GC content as the simplest possible explanation (see tabulated values in Figure 3A,B), we examined whether computationally predicted variations in the mRNA folding stability near the 5’-end are associated with changes in the mRNA levels of synonymous variants, as reported in previous high-throughput studies (Goodman et al., 2013) (Kudla et al., 2009). Although there is no unique definition of the folding stability of an mRNA transcript, a common approach for examining sequence-specific effects near the 5’-end is to compute the folding free energy of the first *L* bases of the transcript, Δ*G*_1,*L*_ (Bentele et al., 2013; Goodman et al., 2013; Kudla et al., 2009). Physically, this quantity represents the free-energy difference between a random coil and the ensemble of all possible base-paired configurations, assuming that the 5’-end does not interact with bases beyond the *L*th position.

Here we take a similar approach by computing a folding free energy that captures the effects of both local and non-local base-pairing. We first hypothesized that the folding of only a portion of an mRNA transcript near the 5’-end is important. Thus, to account for all possible interactions between the first *L* bases and any mutations throughout a transcript, we define

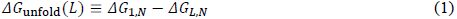

where Δ*G*_1,*N*_ is the folding free energy of the entire transcript and Δ*G*_*L,N*_ includes all contributions from helices that do not involve the first *L* bases (Figure 4A). Equivalently, Δ*G*_unfold_ is equal to the sum of Δ*G*_1,*L*_ and Δ*G*_non-local_, the free-energy difference due to “non-local” contacts between the first *L* bases and bases located farther downstream. Using our complete data set of pBAD constructs, including constructs designed specifically to vary the folding free energy (‘VarE’ sequences in Table S1), we compared 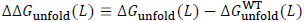 to the log-ratio of the mutant and WT mRNA levels (see Methods). We obtained a maximal Pearson correlation coefficient of 0. 48 (*p*= 4. 5 × 10^−5^) when we chose *L* corresponding to the end of the sixth codon (Figure S1A,B), which suggests that secondary structure formation involving bases in a region stretching from the beginning of the 5’-UTR through the first ~18 bases of the coding region is most important for determining the intracellular mRNA levels; in support of this conclusion, we also found that excluding the 5’-UTR from these secondary-structure calculations results in substantially weaker correlations (Figure S1C,D). Nevertheless, we note that mutations beyond the first six codons can interact with this crucial region by forming intra-molecular secondary structures. With this choice of *L*, the predicted typical magnitude of Δ*G*_unfold_ is a physically reasonable 10*kT*. Calculating this correlation separately for constructs with mutations only in the N-terminal region (Figure 4B) and mutations elsewhere (Figure 4C,D) indicates that synonymous substitutions within the first 20 codons are primarily responsible for the strong overall correlation. Therefore, despite the inherent limitations of current secondary-structure prediction algorithms, we concluded that secondary structure formation near the 5’-UTR plays a central role in dictating the intracellular mRNA abundances, especially when the synonymous substitutions are located near the N-terminus.

**Figure 4:**
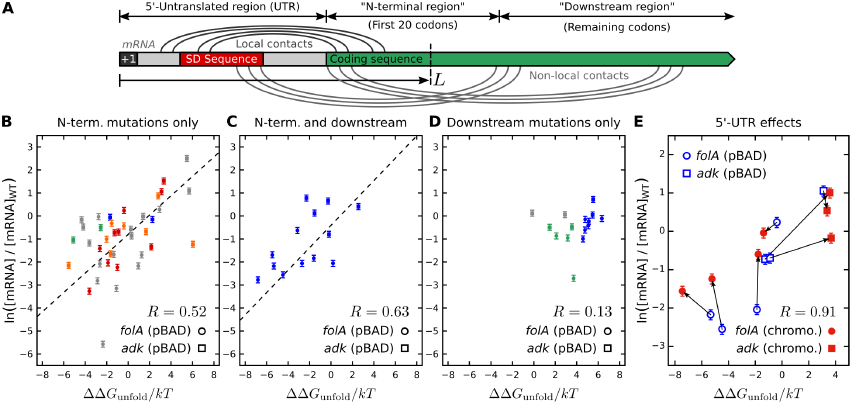
Δ*G*_unfold_ captures the effects of synonymous substitutions in the N-terminal region as well as the effect of the 5’-UTR. (A) A schematic representation of local contacts involving only the first LL bases and non-local interactions. The 5’-UTR, the N-terminal and downstream regions of the sequence are also defined. The log-ratio mRNA levels correlate well with ΔΔ*G*_unfold_, with LL corresponding to the sixth codon, for sequences that (B) contain N-terminal mutations only or (C) contain mutations both in the N-terminal and downstream regions. In (D), this correlation is poor for constructs that only contain downstream mutations. The constructs in panels (B-D) (see Table S1) are color-coded as follows: synonymous replacements of the first rare codon (red); synonymous mutations adjacent to the first rare codon, keeping the latter optimized (orange); systematic mutations of other rare codons (blue); control sequences and other sequences designed to vary Δ*G*_unfold_ (gray); and constructs that incorporate a synonymous anti-SD sequence (green). (E) Relative intracellular mRNA levels for select synonymous mutations engineered on the chromosomal copy of *folA* and *adk* genes in the MG1655 strain (filled symbols) are compared with those on the pBAD plasmid (empty symbols). Identical mutations on the two different backgrounds are connected by arrows, showing that the effects of synonymous substitutions are dependent on the 5’-UTR sequence. For panels (B-E), circles represent *folA* constructs while squares represent *adk* constructs.

### Chromosomal incorporation reveals a strong dependence on the 5’-UTR

These observations imply that mutations in the 5’-untranslated region (5’-UTR) may substantially alter the effects of synonymous substitutions in the coding region. To test this prediction, we introduced select synonymous substitutions in the chromosomal copies of *folA* and *adk* using genome editing. As expected, both ΔΔ*G*_unfold_ and the relative mRNA levels were altered when moving to the endogenous chromosomal promoter and UTR from the pBAD system (Figure 4E). To compare the effects of synonymous substitutions in these different systems and to control for differences in ribosome binding site strengths, we have normalized the mRNA levels with respect to the WT coding sequence on each promoter. In their endogenous contexts, we find that the log-ratio mRNA levels of both genes maintain a strong correlation with ΔΔ*G*_unfold_ (Pearson *R* = 0. 91, *p*= 4. 9 × 10^−3^). Furthermore, as predicted by ΔΔ*G*_unfold_,single synonymous substitutions in two *adk* constructs (6TTG and 6TTA) that have deleterious effects on the mRNA levels when expressed using the pBAD system turn out to have negligible, or even beneficial, effects when chromosomally incorporated.

### Interactions between the Shine–Dalgarno and coding sequences reduce mRNA levels

To better understand the role of mRNA folding stability, we examined the predicted secondary structures of all our constructs with mutations in the 20-codon N-terminal region. Following previous studies (Goodman et al., 2013; Voges et al., 2004), we calculated the probability that a given base is not paired in a transcript’s equilibrium ensemble of secondary structures, *p*_unbound_. Here we analyzed the correlation between the log-ratio of the pBAD mRNA levels and the log-ratio of *p*_nubound_ for mutant and WT transcripts truncated after the first 20 codons, although our results do not change qualitatively when we perform this calculation with full-length transcripts. Using a bootstrap analysis to account for outliers (see Methods), we observed a multi-modal distribution of *p*-values within ~20 bases of the start codon (Figure 5A). Interestingly, the sole local minimum in this distribution that does not correspond to the location of our coding-sequence mutations (i.e., away from codon 3 in *folA* and codon 6 in *adk*) occurs in the middle of the Shine–Dalgarno (SD) sequence, which is part of the pBAD 5’-UTR. In particular, at position −10, we find a moderate, yet significant, correlation (Pearson *R* = 0. 53, *p* = 4. 8 × 10^−4^) between the variations in the measured mRNA levels and changes in *p*_nubound_ (Figure 5B, large symbols).

**Figure 5:**
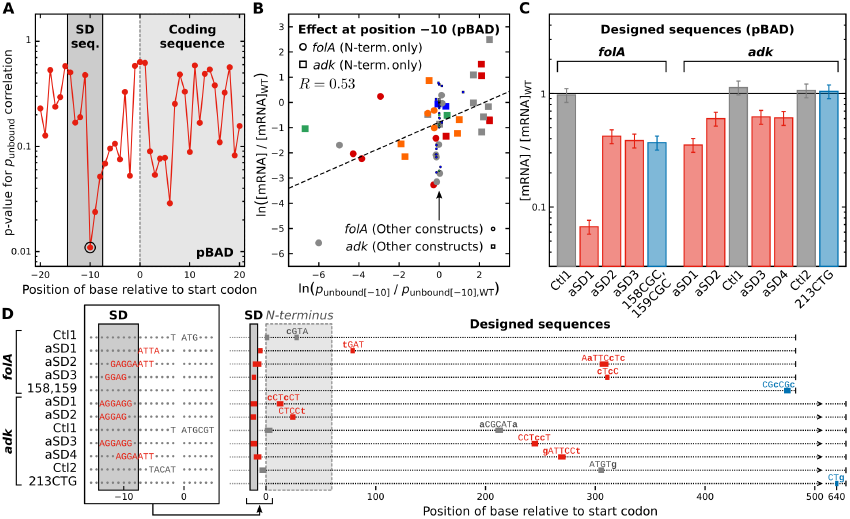
Synonymous substitutions result in occlusion of the Shine–Dalgarno (SD) sequence. (A) The statistical significance of the correlation between the log-ratio mRNA levels and the log-ratio of *p*_unbound_, calculated using transcripts truncated after codon 20, is maximal for the base at position −10, which is located within the SD sequence. These calculations were performed using the subset of constructs with substitutions only in the 20-codon N-terminal region, and the p-values are determined from a bootstrap analysis of these data. (B) By contrast, constructs with substitutions elsewhere in the coding region (small points shown by the arrow) have no dependence on *p*_unbound._ The constructs in panel (B) are color-coded as in Figure 4B-D. (C) Synonymous sequences that incorporate designed anti-SD sequences at different positions in the coding region (red bars) show a substantial drop (40-60%) in mRNA levels on the pBAD system, while control sequences with synonymous mutations that do not contain anti-SD sequences (gray bars) do not exhibit decreases in mRNA levels. Blue bars indicate sequences that were part of our codon optimized mutant collection (Table S1) and contain synonymous substitutions far downstream in the coding region; *folA* 158,159 is partially complementary to the SD sequence. (D) A schematic representation of the designed anti-SD sequences. For each sequence, two subsequences are shown by colored boxes: a target sequence near the 5’-end and an exactly complementary downstream mutant sequence. The mutant bases in the downstream subsequence are shown in lowercase. The 5’-UTR and part of the N-terminus are enlarged in the left panel for clarity, and the color codes are the same as in panel (C). In the case of the anti-SD constructs, the target sequence overlaps with or is immediately adjacent to the SD sequence, while for the control sequences, the target sequence is far from the SD region. This indicates that anti-SD sequences decrease the mRNA abundances irrespective of their positions within the coding region.

This analysis of predicted equilibrium secondary structures suggests that, for synonymous variants with substitutions close to the 5’-end, interactions with the SD sequence are primarily responsible for the correlation between the mRNA levels and ΔΔ*G*_unfold_. However, this approach fails for constructs with substitutions beyond the first 20 codons, where *p*_unbound_ is predicted to be nearly constant but the mRNA levels vary substantially (Figure 5B, small symbols). This observation is not surprising, considering that the predicted secondary structures are dominated by local contacts between bases that are less than ~30 nucleotides apart in the sequence (Figure S2). It is thus apparent that, while equilibrium secondary-structure calculations provide useful information regarding local interactions with the 5’-UTR, purely equilibrium calculations are insufficient for predicting the effects of downstream substitutions.

### Designed downstream synonymous anti-SD mutations reduce mRNA levels

To test the prediction that specific base-pairing with the SD sequence reduces the intracellular mRNA levels, we next engineered synonymous variants containing exactly complementary “anti-SD” sequences throughout the gene. We first designed synonymous variants of *adk* (“aSD1” and “aSD2”) that incorporate an anti-SD sequence within the N-terminal region, while retaining all native rare codons, and observed three-fold and two-fold reductions in mRNA abundance, respectively (Figure 5C). These constructs directly support the hypothesis that intra-molecular base-pairing with the SD sequence is a key determinant of the mRNA levels. We then reasoned that similar effects might occur for anti-SD sequences located throughout the coding region. Because an RNA molecule of length *L* reaches its equilibrium structure at a rate proportional to *L*^−*b*^, where *b* > 1 (Galzitskaya and Finkelstein, 1998), the folding time can far exceed the transcription time, which scales linearly with the transcript length. It is therefore possible that transient contacts between the 5’-UTR and the coding region might sequester the SD sequence while the RNA molecule is out of equilibrium and folding co-transcriptionally; however, such interactions are unlikely to contribute significantly to equilibrium secondary-structure predictions, which are dominated by local contacts.

Because our original data set contained relatively few constructs with far-downstream mutations, we created a larger set of synonymous anti-SD variants (“aSD”), in which anti-SD synonymous sequences, designed to form exactly complementary helices that overlap with or are immediately adjacent to the SD region, were “grafted” far-downstream (Figure 5D). In addition, we designed control variants (“Ctl”) containing sequences that are complementary to portions of the transcript near the beginning of the coding region but away from the SD sequence. As an extra check on the intended complementarity of the mutated regions, we ensured that the designed helices in both the anti-SD constructs and the negative controls are sufficiently stable to decrease ΔΔ*G*_unfold_ by at least 2*kT*. In line with our expectations, we observed a 40-60% reduction in the mRNA levels in pBAD constructs where the grafted anti-SD sequence is far downstream (up to ~300 bases from the SD sequence), while the intracellular mRNA levels of the negative controls were unchanged. We also observed a substantial decrease in mRNA levels for one *folA* construct (‘158CGC+159CGC’), which is partially complementary to the SD region. By contrast, eliminating the last rare codon of *adk* (‘213CTG’) has no effect. These observations therefore demonstrate that interactions with the SD sequence affect the mRNA levels, regardless of the location of the mutations within the transcript. The effects of far-downstream anti-SD sequences in particular highlight the potential for transient, non-equilibrium secondary structures, and not just minimum-free-energy secondary structures, to sequester the SD sequence during mRNA transcription and folding. However, we note that the effects of downstream mutations are less dramatic than those of N-terminal substitutions, as expected for a co-transcriptional folding process.

### Role of co-translational transcription in controlling mRNA levels: Insights from the T7 system

The apparent importance of secondary-structure formation involving the 5’-UTR suggests that variations in the efficiency of translation initiation (de Smit and Van Duin, 1990; Hall et al., 1982), specifically via intra-molecular base-pairing to the SD sequence, lead to changes in the intracellular mRNA abundance. However, two distinct mechanisms could potentially be responsible for these effects. First, the rate of mRNA degradation may increase in the absence of translating ribosomes that would otherwise provide steric protection from nuclease activity (Mackie, 2013; Steege, 2000). Second, since the transcription termination factor Rho holds on to the nascent mRNA during transcription by *E. coli* RNA polymerase, decreasing the rate of ribosome attachment can increase the probability of Rho-dependent transcription termination (Adhya and Gottesman, 1978; Richardson, 1991; Stanssens et al., 1986). To distinguish between these two possibilities, we sought to decouple the processes of transcription and translation through the use of an alternative RNA polymerase. We quantified *in vivo* mRNA and protein levels for selected *folA* constructs cloned under the promoter for T7 RNA polymerase (T7 RNAP), which neither shares subunits with *E. coli* ribosomes nor associates with the termination factor Rho (McGary and Nudler, 2013). Moreover, T7 RNAP travels about 8 times faster than ribosomes on the same mRNA transcript, thereby greatly reducing the probability of co-translational transcription (Iost et al., 1992). The 5’-UTRs of the T7 and pBAD systems are also sufficiently similar to compare the effects of synonymous codons on transcription and translation initiation (Figure S3).

In sharp contrast with the pBAD/Ec RNAP system, we find that the intracellular mRNA levels are largely insensitive to codon substitutions when the *folA* codon variants are cloned under the T7 promoter (Figure 6A). Since transcription is independent of translation in this system, this experiment indicates that the absence of translating ribosomes does not increase the rate of mRNA degradation. However, the variations in soluble protein levels are comparable regardless of the polymerase (Figure 6B), suggesting that similar effects of synonymous mutations on translation initiation alter the efficiency of ribosome attachment whether translation occurs co- or post-transcriptionally. One exception is the fully optimized *folA* sequence (‘MutAll’), which exhibited elevated mRNA levels with T7 RNAP but extremely low mRNA levels with Ec RNAP (Figure 2A). To determine whether MutAll is inherently produced in larger amounts than the WT under the T7 promoter, we carried out *in vitro* transcription of the WT and MutAll sequences with T7 RNAP in the absence of nucleases and found that the MutAll mRNA levels were in fact marginally lower than those of the WT *folA* coding sequence (Figure 6C). We therefore speculate that the high *in vivo* mRNA levels for this construct might be due to a reduced mRNA degradation rate, perhaps by virtue of very stable mRNA secondary structures. Overall, we conclude that in a system where transcription and translation are decoupled (T7 RNAP), inefficient translation-initiation only affects protein levels, while in a coupled system (Ec RNAP), translation-initiation affects both mRNA and protein levels, presumably as a result of premature transcription termination.

**Figure 6:**
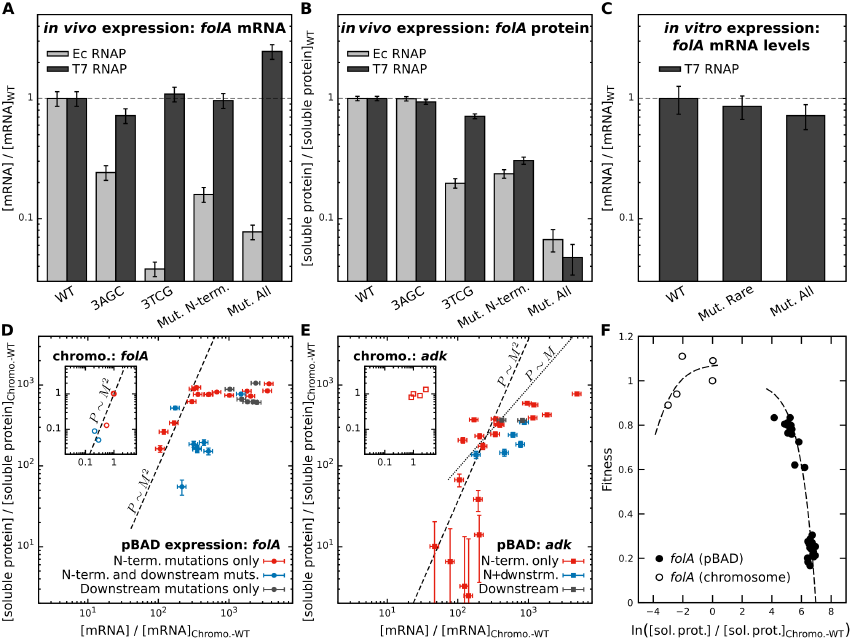
A mechanism of co-translational mRNA transcription explains sequence-specific variations in mRNA levels, protein abundance and cellular fitness. Comparison of (A) relative mRNA and (B) relative soluble protein levels for selected *folA* mutants when expressed under the bacteriophage T7 promoter and under the pBAD promoter. The constructs 3TCG, MutN-term, and 3AGC, which all showed substantial drops in mRNA levels in the pBAD system, have mRNA levels equal to the WT sequence in the T7 system, while MutAll results in a large increase in mRNA but produces very little protein. The trends in protein levels are similar across both the systems. (C) *In vitro* transcription by T7 RNAP. The WT, MutAll and MutRare variants of the *folA* gene were fused to the T7 promoter, and transcription was carried out *in vitro* using T7 RNA polymerase. The amount of transcribed mRNA was quantified using qPCR, and mRNA levels were normalized with respect to the WT. Soluble protein abundances (over-expression in the pBAD system relative to the chromosomal levels, see Table S1) are shown as a function of their mRNA abundances for (D) *folA* and (E) *adk*; note that this normalization hides the fact that *adk* is endogenously expressed at much higher levels than *folA*. The dashed lines illustrate a quadratic relationship between the protein, pp, and mRNA abundances, *M*, at low to moderate expression levels, and the soluble protein abundances appear to plateau relative to the intracellular mRNA abundances at the highest expression levels. The insets show equivalent plots for synonymous variants expressed on the chromosome. (F) Variations in soluble protein abundances arising from synonymous substitutions in *folA* predict cellular fitness (i. e., the exponential-phase growth rate normalized by the WT growth rate, either without the plasmid, for chromosomal incorporations, or at zero arabinose concentration, for the pBAD constructs). The under-expression arm exhibits a Michaelis–Menten-like behavior, while the over-expression arm has a linear dependence on the soluble protein abundance.

### Soluble protein levels vary non-linearly with changes in mRNA levels

We next examined the extent to which protein abundances can be predicted by measuring Ec RNAP-transcribed mRNA levels. We intuitively expected to find a linear relationship between these quantities, in which an increase in the steady-state mRNA levels would lead to a proportional increase in protein production. Yet surprisingly, we instead observed a highly non-linear relationship between the protein and mRNA levels for both genes in both the endogenous and pBAD contexts (Figure 6D,E). In particular, we found an approximately quadratic relationship between the protein and mRNA levels for constructs with low mRNA levels relative to the WT sequence (dashed lines in Figure 6D,E); by contrast, the expected linear relationship (dotted line in Figure 6E) clearly results in a poor fit. Furthermore, at high mRNA levels relative to the WT sequence, the protein levels tend to plateau for both genes. To investigate this latter behavior, we performed a control experiment in which the induction level of WT *folA* on the pBAD plasmid was titrated over three orders of magnitude (Figure S4). This experiment resulted in a similar plateau in the protein levels, indicating that this behavior is not a consequence of the synonymous substitutions specifically, but rather reflects a generic resource limitation at very high over-expression (≳1000-fold over the chromosomal level). Lastly, we found that not all constructs follow the same overall trend, as some pBAD constructs with downstream mutations (blue points in Figure 6D,E) exhibit lower protein levels than most pBAD constructs with similar mRNA levels but N-terminal substitutions only (red points in Figure 6D,E).

To rationalize these unexpected findings, we developed a kinetic model based on the insights that we obtained by comparing the pBAD and T7 *in vivo* transcription systems. We first hypothesized that the non-linear relationship between protein and mRNA levels is due to coupled transcription and translation. To this end, we assumed that ribosomes and Rho termination factors compete for association with nascent mRNA chains, such that transcription and translation both rely on the binding of one of a finite supply of ribosomes. We also assumed that the mRNA transcription and ribosome attachment rates are both proportional to the accessibility of the 5’-UTR, as predicted by Δ*G*_unfold_. In this model, proteins can then be produced either co-transcriptionally by the leading ribosome that ensures complete transcription, or post-transcriptionally by ribosomes that subsequently attach to the transcribed mRNA molecules (Bakshi et al., 2012). With these assumptions, we show in the SI Text that the steady-state rate of protein production has a quadratic dependence on the steady-state mRNA abundance under conditions where translation initiation is inefficient due to the inaccessibility of the SD sequence. In this regime, the reduced rate of translation initiation greatly increases the probability of premature transcription termination, such that protein production is dominated by the post-transcriptional translation of a relatively low concentration of transcripts. This model therefore supports the notion that the non-linearity of the observed relationship arises, at least in part, from the co-transcriptional nature of translation in *E. coli*. In addition, this approach suggests that both the plateau in protein expression and deviations from the general trend exemplified by constructs with N-terminal mutations may arise from variations in mRNA degradation rates, due either to a generic exhaustion of the available nuclease activity or to sequence-specific effects, as seen in the case of MutAll (Figure 6A).

We then examined whether an understanding of the mRNA and protein levels is sufficient to predict any fitness effects associated with synonymous substitutions. As shown in Figure 1, chromosomal incorporations of mutated *folA* sequences reduce the exponential-phase growth rate of *E. coli*; here we show that synonymous mutations in over-expressed DHFR also result in substantial fitness effects (Figure 6F). Despite the potential for synonymous substitutions to affect cellular fitness via multiple mechanisms, including effects due to co-translational protein folding and aggregation, we found that in the case of *folA*, variations in the exponential-phase growth rate are well explained by the soluble protein levels alone. Protein-level variations in both the under and over-expression regimes lead to fitness effects that are consistent with our previous study of WT DHFR expression levels (Bhattacharyya et al., 2016). In particular, in the DHFR under-expression regime, the dependence of the cellular growth rate on DHFR abundance is Michaelis–Menten-like (Figure 6F, empty circles), while in the over-expression regime, this dependence is linear (Figure 6F, filled circles). Nevertheless, we emphasize that, unlike our prior study, under-expression is caused solely by synonymous substitutions within the chromosomal copy of *folA*, while variations in over-expressed protein levels are caused by synonymous substitutions at a fixed inducer concentration. Lastly, the strong linear relationship between variations in the total and soluble protein abundances that we obtain for both DHFR and ADK (Figure S5), both on the plasmid and on the chromosome, indicate that aggregation does not measurably increase upon over-expression.

### A genome-wide analysis reveals selection to avoid Shine–Dalgarno mis-interactions

Motivated by the clear fitness effects of synonymous mutations in our experiments on *folA*, we undertook a statistical analysis of the *E. coli* transcriptome to evaluate the generality of our conclusions. We hypothesized that widespread selection against synonymous sequences that occlude the SD region might lead to detectable differences in the folding free energies and base-pairing patterns between the extant transcripts and synonymous alternatives for a large number of *E. coli* genes. Using the final 20 bases of the 5’-UTR and the entire coding sequence, we first found the mean Δ*G*_unfold_ (with *L* corresponding to the fifth codon) of the WT sequences to be approximately −12KKKK (Figure 7A). We then constructed a control data set by considering all possible synonymous variants of the first five codons of each *E. coli* gene. Assuming that the synonymous codons are selected independently and with equal frequencies, the most probable Δ*G*_unfold_ distribution is shifted to the left of the WT distribution (Fig. 7A). This observation is consistent with selection against strong interactions between the SD and coding sequences. To make a quantitative comparison between these histograms, we computed the probability, within the control distribution, of observing no more than the actual number of genes with Δ*G*_unfold_ less than a specified threshold. The resulting *p*-values indicate that the control and WT distributions differ most significantly with respect to the number of genes that have Δ*G*_unfold_ ≲ −14*kT* (Figure 7B, red curve). This conclusion is unchanged if we instead bias the selection of synonymous codons in accordance with the empirical genome-wide codon-usage frequencies within the first five coding positions (Figure 7C, blue curve). This analysis demonstrates that the sequence context, and not simply the relative usage of each codon, contributes to the difference between the WT and the control Δ*G*_unfold_ distributions.

**Figure 7:**
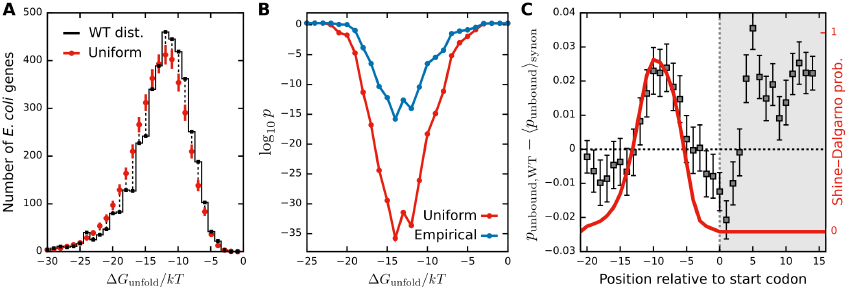
A statistical analysis of E. coli N-terminal codon bias provides evidence for selection against SD mis-interactions. (A) A comparison of the genome-wide distributions of Δ*G*_unfold_ for the wild-type (WT) and alternative synonymous sequences, assuming that all synonymous codons are used with equal frequencies. The first 15 bases of the coding region are mutated, and we have used *L* corresponding to the fifth codon in these calculations. (B) p-values (see text) assess the statistical significance of the difference between the WT and control distributions, assuming that synonymous codons are selected either uniformly or according to the empirical genome-wide usage within the first five codons. (C) The average difference in the equilibrium base-pairing probability between the WT and the mean of the uniformly weighted synonymous sequences for each gene. The most significant decrease in secondary-structure formation within the 5’-UTR (unshaded region) coincides with the most probable location of the SD sequence (red curve); see also Figure S6.

We then sought to determine whether *E. coli* genes are optimized to avoid base-pairing with the SD sequence. Based on our analysis of *folA* and *adk* mRNA levels, we reasoned that base-pairing near the 5’-end of a transcript can be predicted reasonably well using equilibrium secondary-structure calculations. We therefore calculated *p*_unbound_ for the final 20 bases of the 5’-UTR and the first 5 codons for all *E. coli* genes and their synonymous variants. By looking at the difference between *p*_unbound_ in each WT sequence and the mean of the uniformly weighted synonymous variants, we detected a statistically significant reduction in base-pairing within the 5’-UTR of the WT genes (Figure 7C). We then located the SD sequence for each gene by minimizing the hybridization free energy between a 7-base sliding window and the *E. coli* rRNA anti-SD sequence, verifying that the resulting distribution of SD-sequence locations agrees with previous experimental studies of SD-sequence placement (Chen et al., 1994). Remarkably, the greatest reduction in predicted base-pairing in the 5’-UTRs of the WT transcripts coincides precisely with the most likely location of the SD sequence, indicating that the reduction in mRNA secondary structure near the 5’-end is driven specifically by an avoidance of interactions that tend to occlude the SD sequence. Since helical secondary structures are formed by base-pairing between two disjoint subsequences in an RNA molecule, it is not surprising that we also observe a commensurate reduction in WT base-pairing in the 15 bases that we specifically mutated within the coding region. To this end, we verified that this result is qualitatively unchanged by increasing the number of coding-region bases considered in this calculation, as we observe no statistically significant difference in *p*_unbound_ between the WT transcripts and the synonymous ensembles for bases beyond the mutated five codons (Figure S6). Thus, by directly implicating the SD sequence, this statistical analysis lends additional support to our proposed SD-occlusion mechanism.

## Discussion

Codon usage bias has been a fascinating topic of research for over 30 years (Brule and Grayhack, 2017; Chaney and Clark, 2015; Plotkin and Kudla, 2011). While uneven codon usage has been firmly established, the biological causes and consequences of non-random synonymous-codon selection remain largely unknown. Identifying specific mechanisms is challenging, since codon usage can affect all stages of protein biosynthesis, from transcription and translation to co-translational protein folding and potentially protein-complex assembly. Variations in codon usage near the N-terminus, where the relative frequencies of synonymous codons tend to be most dissimilar from the remainder of the coding sequence (Clarke and Clark, 2010), are particularly intriguing, as substitutions near the beginning of a gene are unlikely to affect co-translational folding or assembly. Instead, it is widely believed that this N-terminal codon bias arises in some way from the behavior of mRNA transcripts.

Previous experimental approaches to assess the effects of N-terminal synonymous substitutions have used libraries of synonymous variants of several endogenous and non-endogenous genes expressed in *E. coli* (Boel et al., 2016; Goodman et al., 2013; Kudla et al., 2009; Welch et al., 2009). These studies have provided a plethora of data for statistical analyses that can be used to assess the primary determinants of codon-level effects on protein expression. All of these studies showed large variations in protein expression, often reaching several orders of magnitude, as a result of synonymous substitutions near the N-terminus. Perhaps surprisingly, a similar or greater variability was sometimes observed for mRNA levels, suggesting a link between synonymous codon usage and mRNA turnover. Linear regression analyses pointed to multiple relevant factors, including mRNA secondary structure stability, deleterious effects of specific codons, and several other signatures that were shown to predict protein expression in gene-design applications (Boel et al., 2016; Welch et al., 2009). However, these approaches have not provided a complete understanding of the physical mechanisms and evolutionary factors responsible for the observed biases in codon usage.

To address these challenges, we chose an alternative, bottom-up approach where we explored possible mechanisms by making one codon substitution at a time. By designing synonymous variants of two endogenous *E. coli* genes, we tested hypotheses regarding the relative importance of codon rarity, the context of rare codons within a sequence, and multiple predictors of mRNA folding stability. In line with earlier studies, we observed dramatic variations in both mRNA and protein expression levels. However, our systematic analysis also revealed highly position-specific effects of synonymous substitutions on mRNA and protein levels. We found that synonymous replacements of specific rare codons near the N-terminus, and in particular the first rare codon (encoding Ser3 in *folA* and Leu6 in *adk*), tend to have the greatest effects. Nevertheless, codon rarity *per se* is not a determining factor, since replacing WT rare codons with either optimal or alternative rare codons had similar detrimental effects. This observation argues against the importance of translational pauses near the 5’-end (Eriksen et al., 2017; Mitarai et al., 2008; Pedersen et al., 2011).

Our results are instead consistent with the hypothesis that the 5’-end of a transcript should be free of secondary structure to promote efficient translation initiation (Eriksen et al., 2017; Ringquist et al., 1992). Yet in contrast to previous studies that proposed a relatively non-specific mRNA folding stability for the first ~120 bases of the transcript (Boel et al., 2016; Goodman et al., 2013; Kudla et al., 2009), our analysis indicates that the region of the transcript that should be free of secondary structures extends only from the 5’-UTR through the first ~6 codons. In particular, we have shown that the observed variations in mRNA levels correlate most strongly with the presence of secondary structures involving the SD sequence. Nevertheless, we emphasize that because the formation of helical secondary structures in RNA molecules always involve a *pair* of complementary sequences, mutations that sequester the SD sequence may be located downstream of the sixth codon. When such coding-region substitutions are relatively close to the N-terminus, the relevant interactions with the 5’-UTR can be predicted using equilibrium secondary-structure calculations, as demonstrated by our analysis of *p*_unbound_ in the first ~20 codons. However, mutations that occur farther downstream can also lead to transient interactions with the 5’-UTR during the process of co-transcriptional folding. This hypothesis is supported by the behavior of synonymous *folA* and *adk* constructs that incorporate anti-SD sequences while retaining all native rare codons. Unlike control sequences that lack such motifs, our anti-SD constructs exhibited substantially reduced mRNA levels, regardless of whether the anti-SD motif was engineered in the N-terminal region or as far as 300 bases downstream. These effects further establish the importance of the SD sequence and highlight the potential for non-equilibrium, transient secondary structures to alter the efficiency of protein production when transcription, mRNA folding, and translation all occur on similar timescales *in vivo*. To the best of our knowledge, we believe that this is the first study that shows the effect of far-downstream mutations and non-equilibrium secondary structures on mRNA levels; however our treatment of these non-equilibrium effects is only approximate, and further theoretical analysis is needed to understand this behavior completely.

A major conclusion from our study is that the effect of synonymous codons on mRNA and protein levels depends on the RNA polymerase that transcribes the mRNA. In fact, the same synonymous mutations can either increase or decrease mRNA levels depending on whether *E. coli* RNAP or T7 RNAP is used. This difference is due to the co-transcriptional nature of translation in *E. coli*, while the non-endogenous T7 RNAP is free from these effects. Earlier studies that used EcRNAP (Goodman et al., 2013; Kudla et al., 2009) observed decreases in mRNA levels upon codon optimization, while those that used the T7 system (Boel et al., 2016) observed an increase in mRNAs for optimized genes. However, it was difficult to resolve this controversy because these studies used different sets of gene constructs. Our study is unique in that we examined the same constructs using both transcription systems and thus obtained a clearer understanding of the role of co-translational transcription in dictating codon usage. Our study also implicates an interesting role of the transcription termination factor Rho in controlling mRNA levels, although further work is required to address this in detail. Moreover, our results obtained in the T7 system may have direct implications for codon usage bias in eukaryotes, where transcription and translation occur in separate cellular compartments.

Finally, we linked the effects of synonymous substitutions to changes in cellular fitness, both in the specific case of *folA* and, more generally, for the *E. coli* genome as a whole. Comparing the expression of synonymous variants on the chromosome versus the plasmid, we found that the WT *folA* sequence appears to be well optimized to maximize protein production under its endogenous promoter. Then, to corroborate our mechanistic hypothesis, we used a statistical analysis of the *E. coli* genome to show that extant mRNA transcripts specifically avoid SD-sequence base-pairing relative to alternative synonymous sequences. Notably, this difference cannot be explained simply by the relative frequencies of synonymous codons near the N-terminus, which again emphasizes the importance of sequence context in determining the interactions between the coding and SD sequences. These results are compatible with the conclusions of (Bentele et al., 2013), but go further by specifically implicating the avoidance of SD sequence interactions, as opposed to more generic secondary-structure formation in the vicinity of the 5’-end. This analysis therefore corroborates our experimental conclusions on a much larger scale and indicates that the physical mechanism that we have proposed has strongly influenced the evolution of codon selection near the N-terminus.

In summary, we have used a systematic analysis of synonymous substitutions in two endogenous *E. coli* genes to establish a mechanism of N-terminal codon bias. We have shown that secondary structure formation involving the Shine–Dalgarno sequence is a primary determinant of optimal codon selection due to its crucial function in promoting translation initiation, which modulates both mRNA and protein expression levels through co-transcriptional translation in *E. coli*. Our analysis improves upon earlier studies by validating a specific definition of the folding stability of mRNA transcripts near the 5’-end and revealing the previously unrecognized, yet crucial, role of co-transcriptional mRNA folding. Our model therefore connects “silent” mutations, through RNA folding dynamics, to their effects on cellular fitness, providing a general mechanistic insight into the origin of codon-usage bias.

## Methods

### Definition of rare codons

We consider a codon to be rare if its usage within the most abundant *E. coli* proteins (Ikemura, 1985) is less than 10% of the usage of the amino acid for which it codes.

### Plasmids and strain construction

We cloned synonymous codon variants of *folA* and *adk* genes in the pBAD plasmid between *NdeI* and *XhoI* sites under control of an arabinose inducible promoter. We used a megaprimer based PCR method or an inverse PCR method using partially complementary primers for site-directed mutagenesis to mutate codons on the plasmid. For both methods, KOD Hot Start DNA polymerase was used. The fully optimized ‘MutAll’ sequence and the rare-optimized ‘MutRare’ sequences for both *folA* and *adk* were synthesized as g-block gene fragments from IDT, and cloned in pBAD vector. Expression of pBAD plasmid in all cases was done in *E. coli* BW27783 cells (CGSC #12119).

We made MG1655 strains having synonymous mutations in the chromosomal copies of the *folA* and *adk* genes using a genome editing technique described previously (Bershtein et al., 2012).

### Media conditions

We carried out bacterial growth in M9 minimal salts supplemented with 0. 2% glucose, 1mM MgSO_4_, 0. 1% casamino acids, and 0. 5 μg/ml thiamine (supplemented M9 medium). For BW27783 cells containing the pBAD plasmid, ampicillin was added to a concentration of 100μg/ml during growth. mRNA and protein abundance measurements were carried out using a saturating arabinose concentration of 0. 05%.

### mRNA abundance measurement

To determine *in vivo* mRNA levels, MG1655 strains containing chromosomally engineered mutations or BW27783 cells transformed with pBAD plasmid or BL21 strains transformed with pET24a(+) plasmid were grown overnight in supplemented M9 medium at 37°C. The next day, the culture was diluted 1:100 into fresh M9 medium (containing 0. 05% arabinose for pBAD plasmid or 50μM IPTG for pET24a plasmid) and grown for 4 hours at 37°C. Based on OD_600_ of the cultures, a culture volume corresponding to 5×10^8^ cells was spun down, re-suspended in 1ml of M9 medium, and treated with 2ml of RNAprotect Bacteria reagent (Qiagen), after which the total RNA was extracted using an RNeasy mini kit according to the manufacturer’s instructions. The concentration of the purified RNA samples was estimated using nanodrop. 1μg of the purified total RNA was subjected to reverse transcription to produce cDNA using a QuantiTect Reverse Transcription kit (Qiagen). The levels of *folA* and *adk* genes were estimated by quantitative PCR (qPCR) using 20bp gene specific primers and a QuantiTect SYBR Green PCR kit (Qiagen). For synonymous variants of *folA* gene on the plasmid, chromosomally expressed *adk* gene was used as a reference, while for synonymous variants of *adk* gene on the plasmid, chromosomally expressed *tufA* gene was used as a reference.

### Protein abundance measurement

Protein abundance measurements were done essentially as described in (Bhattacharyya et al., 2017; Bhattacharyya et al., 2016). Briefly, MG1655 strains containing chromosomally engineered mutations or BW27783 cells transformed with pBAD plasmid or BL21 strains transformed with pET24a(+) plasmid were grown in supplemented M9 medium (containing 0. 05% arabinose for pBAD plasmid or 50μM IPTG for pET24a plasmid) for 4 hours at 37°C, chilled on ice for 30 min and spun down. Based on the measured OD600 of the cultures, the pellets were re-suspended in lysis buffer (1×BugBuster (Novagen) and 25units/ml of Benzonase), such that the final OD600 is 2. 0. 10μl of the lysate (before centrifugation) was mixed with 2μl of 50% glycerol+10% SDS, heated at 95°C for 10 minutes, and used for estimation of the total protein abundance. The remaining lysate was centrifuged for 30 minutes at high speed and the supernatant was used for estimation of the soluble protein abundance. Intracellular DHFR and ADK protein amounts in both the total and soluble fractions were determined by SDS-PAGE followed by Western Blot using rabbit anti-DHFR/anti-ADK polyclonal antibodies (custom raised by Pacific Immunology). The intensity of the bands was further normalized by the total protein concentrations obtained using a BCA assay. A representative western blot image is shown in Figure S7.

### Error analysis

The differences between replicates were plotted versus the replicate averages for both mRNA and soluble-protein measurements (Figure S8). Linear fits were performed to estimate the absolute errors, which are constant, and the relative errors, which are proportional to the mean value of each measurement, under the constraint that the absolute error is non-negative. Four *adk* constructs, for which the soluble-protein measurements fall below the absolute error estimate, were excluded from the soluble-protein correlation calculations. The error bars shown in Figures 1, 3, and 4 indicate the estimated standard deviation of the measurements. The error bars in Figure 2 indicate the standard error of the mean of three replicates.

### *In vitro* transcription

The coding regions of WT, MutAll and MutRare versions of *folA* gene were PCR amplified with primers to incorporate a T7 promoter upstream of the coding region, and this linearized DNA was used as a template for *in vitro* transcription using a Maxiscript T7 kit (Thermo Fisher Scientific) using the manufacturer’s instructions. The DNA template was digested with the Turbo DNase included in the kit, after which the mRNA was purified using an RNeasy mini kit, reverse transcribed to generate cDNA, and quantified using qPCR as described above.

### Fitness measurements

Overnight cultures of BW27783 cells containing pBAD plasmid were grown with ampicillin at 37°C while MG1655 strains containing chromosomally engineered mutants of *folA* and *adk* were grown at 30°C. The next day, cultures were diluted 1:100 and grown at 37°C for 12 hours in Bioscreen C system (Growth Curves USA). For BW27783 cells, 0. 05% arabinose was added for induction. OD data were collected at 600nm at 15 min intervals. The resulting growth curves were fit to a bacterial growth model to obtain growth rate and lag time parameters (Zwietering et al., 1990).

### RNA folding calculations

We used NUPACK (Zadeh et al., 2011) for all RNA folding calculations. This program computes the partition function, ZZ = ∑ss exp (−Δ*g*_*ss*_/*kT*), where the sum runs over all secondary structures, excluding pseudo-knots. The free energies of individual secondary structures, Δ*g*_*ss*_, are predicted using experimentally determined RNA base-pairing rules. The folding free energy of a transcript is then defined as Δ*G*_1,*N*_ = −*kT* log *Z*. Unless otherwise noted, the full transcript, from the predicted +1 site to the stop codon, was used. Calculations were performed assuming standard conditions ([Na^+^] = 1 M; *T* = 37°C), with the results reported relative to the thermal energy, *kT*. While these conditions are not physiological, we expect the relative free-energy differences among synonymous sequences to be robust.

The equilibrium probability that a base is unbound is given by *p*_unbound,*i*_ = 1 − Z^−1^ ∑_ss_ **1**_*i,ss*_ exp (−Δ*g*_*ss*_/*kT*), where 1_*i,ss*_ indicates that the base at position *i* is paired in a given secondary structure. To examine the role of specific bases (Figure 5C), we used a bootstrap analysis to account for the effect of outliers on the correlation between the log-ratio mRNA levels and the log-ratio of *p*_unbound,*i*._ To this end, we sampled our synonymous constructs with replacement to create approximately 1,000 alternative datasets with the same length but randomly repeated entries. The bootstrapped *p*-value is then the arithmetic mean of the one-tailed Pearson correlation *p*-values for all generated samples.

## Supplementary figure captions

**Figure S1:**
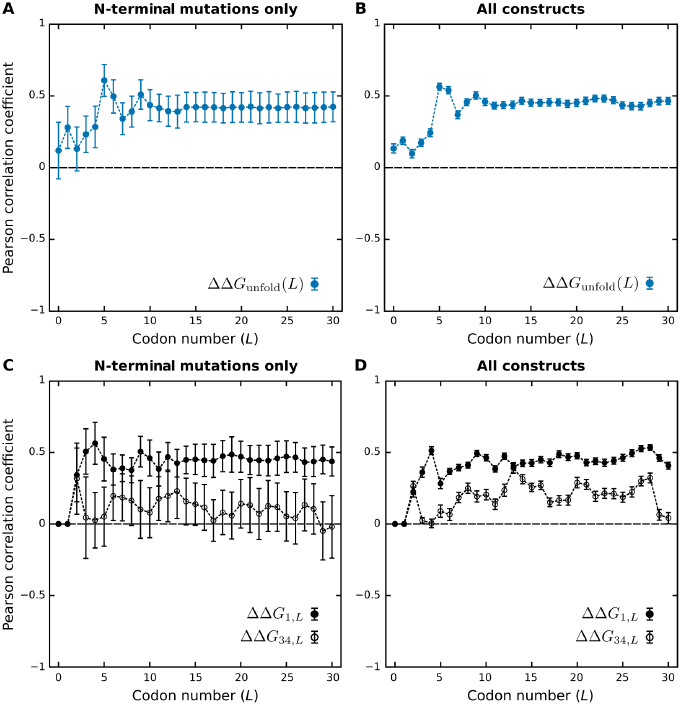
(A-B) The Pearson correlation coefficient, relating the log-ratio mRNA levels and ΔΔ*G*_unfold_, as a function of *L* (See Figure 4A). Error bars and dashed lines represent the standard deviation obtained from bootstrap analyses of these calculations, as explained in the Methods. In (A), only the 31 pBAD constructs with mutations confined to the first 20 codons are included in the correlation calculations, while in (B), all 65 mutated pBAD constructs are considered (See Table S1). In (B), the correlation with ΔΔ*G*_unfold_ is greatest near *L* corresponding to the sixth codon, indicating that the inclusion of additional downstream bases to the region of interest [1, *L* ≈ 51] bases does not improve our ability to predict the effects on mRNA levels. (C-D) The Pearson correlation coefficient obtained by comparing the log-ratio mRNA levels and the folding free energies of transcripts truncated at base LL. The calculations ΔΔ*G*_1,*L*_ and ΔΔ*G*_34,*L*_ respectively include and exclude the 5’-UTR of the pBAD plasmid; the weaker correlations using ΔΔ *G*_34,*L*_ also point to the crucial role of the 5’-UTR. We chose to use ΔΔ*G*_unfold_ in our analysis, as all mutations in the coding region can be considered in the calculations regardless of the choice of *L*; by contrast, ΔΔ*G*_1,*L*_ requires that we exclude mutations beyond a given codon in order to ignore secondary structures that form downstream of the region of interest [1, *L*]. Consequently, our approach using ΔΔ *G*_unfold_ provides more robust evidence of the importance of the first ~50 bases, including the 5’-UTR.

**Figure S2:**
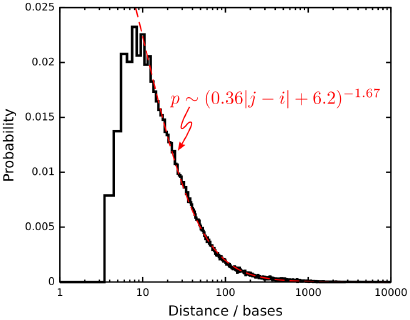
The probability distribution of the distance (with respect to the primary sequence) between paired bases *i* and *j* in the predicted minimum-free-energy secondary structures of *E. coli* mRNA transcripts.

**Figure S3:**
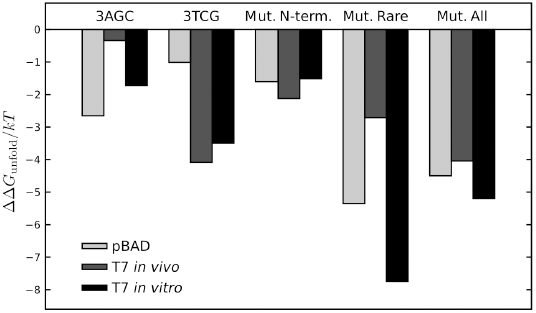
The predicted values of ΔΔ*G*_unfold_, with *L* corresponding to the sixth codon, for selected *folA* constructs on the pBAD plasmid, *in vivo* T7 plasmid, and *in vitro* T7 system. Because the *in vivo* T7 5’-UTR contains the lac repressor binding site and is thus much longer than the 33-base 5’-UTR of the pBAD system, only the final 33 bases were used for these calculations.

**Figure S4:**
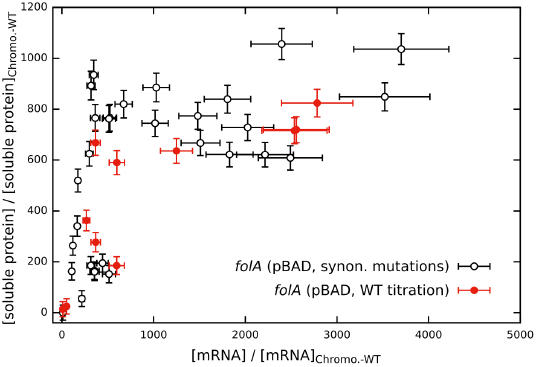
Soluble protein abundances of *folA* synonymous codon variants appear to plateau relative to the intra-cellular mRNA abundances at the highest expression levels. Soluble protein abundances of all variants were measured using 0. 05% arabinose on the pBAD plasmid and normalized with respect to the WT level at this inducer concentration. For comparison, red circles show the soluble abundance of WT DHFR obtained from a range of arabinose concentrations (0. 0000122% to 0. 05%).

**Figure S5:**
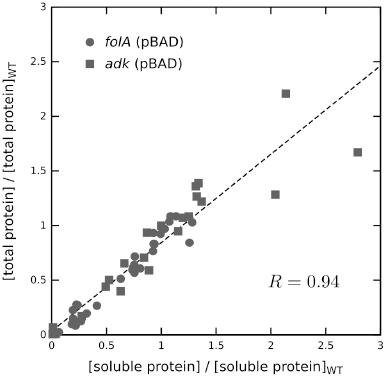
The normalized soluble and total intra-cellular protein abundances exhibit a clear linear dependence. Consequently, the aggregation propensity of both proteins does not appear to increase substantially upon over-expression. We note that [soluble]_WT_/[total]_WT_ ≃ 0. 9 for *adk* and ≃ 0. 95 for *folA*.

**Figure S6:**
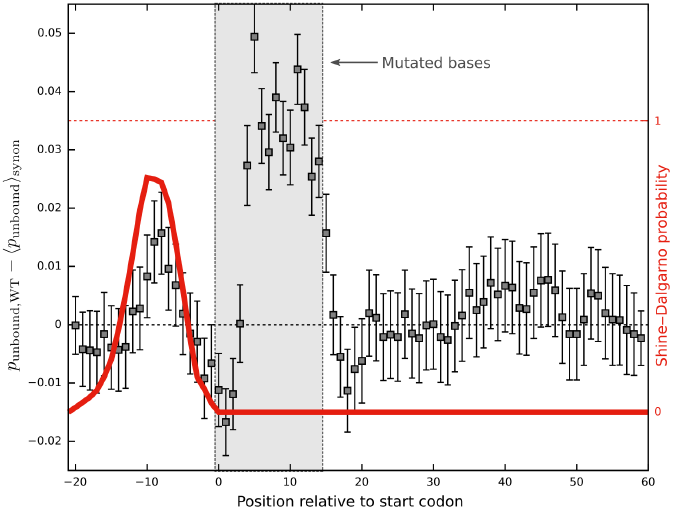
The same calculation as shown in Figure 7C, but using mRNA transcripts truncated after the 20^th^, as opposed to the fifth, codon. As in Figure 7C, only the first five codons were mutated. Aside from those codons that were directly mutated (gray region), the positions where the difference between *p*_unbound,WT_ and the mean *p*_unbound_ for the ensemble of equally weighted synonymous variants is positive and statistically significant (i.e., outside the error bars) are all located within the typical SD region (red curve).

**Figure S7:**
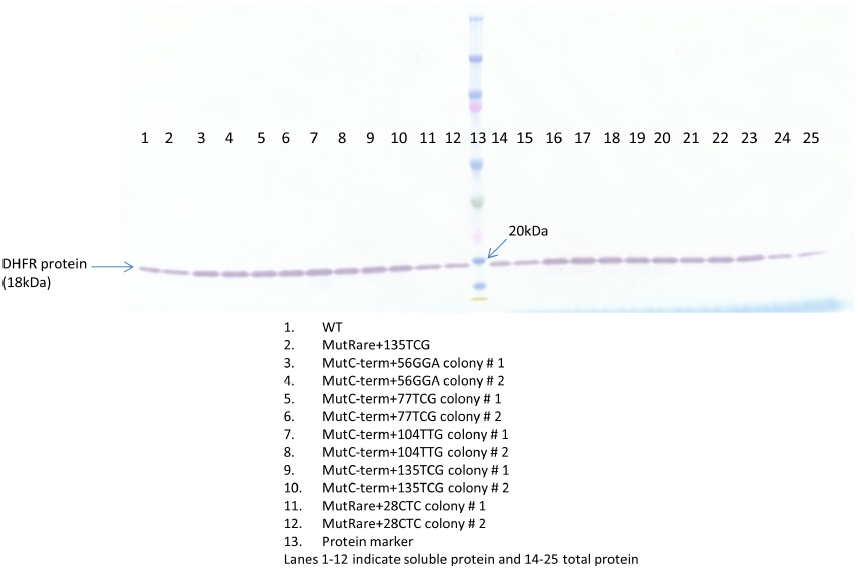
A representative western blot image for different *folA* mutant constructs. The band intensity was quantified using ImageJ software, and further normalized by the total protein concentration of the sample obtained by a BCA assay. The lanes are numbered and described in the figure legend.

**Figure S8:**
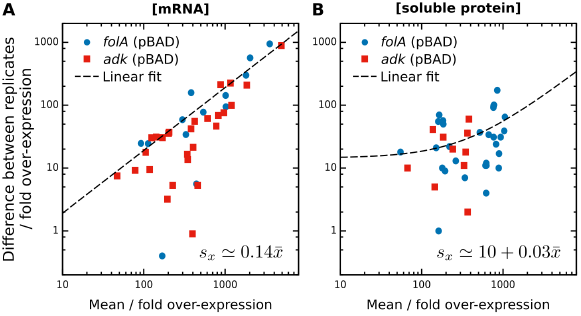
Error analysis for mRNA and soluble-protein levels on the plasmid. Linear fits were used to estimate the typical difference between replicates (see Methods). We then calculated the average standard deviation of each measurement,*s*_*x*_, as a function of the mean of each measurement, 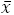 All values are shown here relative to the WT chromosomal expression level.

**SI Text for “Accessibility of the Shine–Dalgarno sequence dictates N-terminal codon bias in *E. coli*”**

Sanchari Bhattacharyya, William M. Jacobs, Bharat V. Adkar, Jin Yan, Wenli Zhang, & Eugene I. Shakhnovich

## Kinetic model of coupled transcription and translation

### A. Kinetic model

We developed a kinetic model of coupled transcription and translation to predict the relationship between the variations in mRNA and protein levels in *E. coli* that result from synonymous substitutions. We assumed that ribosomes and Rho termination factors compete for association with nascent mRNA chains, such that transcription and translation both rely on the binding of one of a finite supply of ribosomes. We then formulated a minimal kinetic model to describe the coupling of the mRNA and protein abundance:

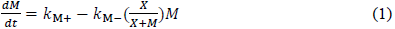

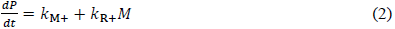

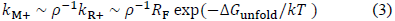

Here, *M* is the total mRNA concentration, *X* is the effective nuclease concentration, *R*_F_ is the concentration of unbound ribosomes, and *dP/dt* is the rate of protein production. The mRNA transcription and ribosome attachment rates, *k*_M+_ and *k*_R+_, are assumed to be proportional to the accessibility of the 5’-UTR, as predicted by Δ*G*_unfold_. However, the transcription rate is decreased by a dimensionless factor *ρ*^−1^, which accounts for premature termination due to competition with Rho. We generally consider the mRNA degradation rate *k*_M−_ to be constant for all transcripts, since our *in vivo* experiments using T7 transcription (Figure 6A,B) indicate that differential mRNA degradation does not play a major role for constructs with N-terminal mutations only; nevertheless, the implications of sequence-dependent mRNA degradation rates are discussed below.

It is straightforward to see that the steady-state mRNA concentration,

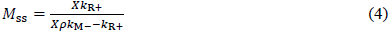

is determined by the balance of the mRNA transcription and degradation rates. Consequently, *M*_*ss*_ is proportional to *k*_R+_, and is thus an exponential function of ΔΔ*G*_unfold_, in the regime where the available concentration of nucleases is high enough to degrade the intracellular mRNA, such that 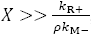, and the degradation rate, *k*_M_, is sequence-independent. As expected, this behavior is consistent with the correlations shown in Figure 4B and Figure S1.

Equation (2) states that proteins are produced either by the first ribosome that is also responsible for ensuring complete transcription, or by ribosomes that subsequently attach to the available mRNA transcripts. The rate of protein production can be written as a function of *M*_*ss*_at steady state,

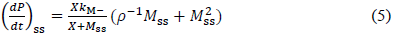

Importantly, the effects of synonymous substitutions on *k*_M+_ and *k*_R+_do not appear explicitly in Equation (5), except through the factor *ρ*^−1^, as discussed below.

### B. Linear and quadratic regimes at low to moderate expression levels

Let us first assume that *M*_*ss*_ is in the linear regime where 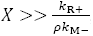. Then Equation (5) predicts that the relationship between the protein-production rate and the steady-state mRNA levels may be either linear or quadratic. If translation initiation is very efficient, such that ribosomes out-compete Rho, then *ρ* is small, and we obtain a linear dependence of the protein levels on the steady-state mRNA levels. This behavior is expected to occur for synonymous sequences that produce mRNA at similar levels to the WT sequence. Furthermore, under these conditions, translation is predicted to occur primarily co-transcriptionally, with the lead ribosome contributing significantly to the total protein production.

If premature transcription termination is instead much more common than translation initiation, such that *ρ* is large, then we expect a quadratic dependence of the protein levels on the steady-state mRNA levels; such behavior is therefore expected at relatively low mRNA levels. In this regime, translation primarily occurs post-transcriptionally, because the initial step of transcription is rarely completed. We note that this quadratic relationship is nevertheless a consequence of the assumed coupling of transcription and translation, which both require ribosome binding. If, instead of Equation (3), we were to assume that the rates *k*_M+_ and *k*_R+_ are independent, then a quadratic relationship would not be predicted by the model.

Evidence for both of these regimes can be seen by comparing the soluble-protein and steady-state mRNA levels for our *folA* and *adk* constructs, both on the chromosome and using the pBAD overexpression system (Figure 6D,E). The quadratic relationship is sketched at low (relative to the WT) mRNA levels by a dashed line. By contrast, we find that the linear relationship that one might expect for decoupled transcription and translation provides an overall poor fit to the data; for example, see the dotted line sketched in Figure 6E.

### C. Plateau in protein production at high expression levels

If mRNA production is so rapid that the available nucleases are overwhelmed, such that 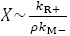, then it is possible for dP/dt to plateau while *M*_*ss*_ increases to very high levels. Obtaining such a plateau from Equation (5) further requires that ρ be small, which is consistent with the regime where the mRNA levels are overexpressed with respect to the WT sequence. In this scenario, we predict that the protein production is still dominated by co-transcriptional translation, but *M*_*ss*_ increases because the concomitantly produced transcripts are not rapidly degraded. The comparison with titrated WT DHFR shown in Figure S4 indicates that this is a generic effect that is not dictated by the transcript sequence itself, but rather originates from a resource limitation in the cell. However, it is unlikely that the limiting resource is the supply of ribosomes, since our experiments comparing *in vivo* transcription using Ec and T7 RNAP (Figure 6A,B) indicate that ribosomes are necessary for both mRNA and protein production, as expressed in Equation (3). We therefore believe that the mechanism of nuclease exhaustion proposed in this model is a more likely explanation for the observed plateau in protein production.

### D. Possible effects of differential mRNA degradation

Finally, we note that the presence of some downstream mutations in the pBAD system appears to increase the mRNA levels relative to constructs with only N-terminal substitutions but otherwise similar protein levels (blue points in Figure 6D,E), as these points fall below the general trend of those constructs with N-terminal substitutions only (red points in Figure 6D,E). This effect may be due to differential mRNA degradation rates arising from synonymous substitutions, as seen in the case of T7-transcribed MutAll in Figure 5A,B. Differential degradation implies that *k*_M_-is sequence-dependent. While variations in *k*_M_-unsurprisingly affect the steady-state mRNA levels, as predicted by Equation (4), Equation (5) also predicts that such variations affect the relationship between (*dP/dt*)_*ss*_ and *M*_*ss*_. This mechanism is thus a plausible explanation for the observed behavior of the MutAll construct. Nevertheless, we emphasize that the modulation of mRNA and protein levels through the effects of translation initiation on transcription, as opposed to differential mRNA degradation, appears to be the dominant mechanism dictating N-terminal codon bias, as we argue in the main text.

